# Sensory learning and inference is impaired in the non-clinical continuum of psychosis: a replication study

**DOI:** 10.1101/296988

**Authors:** Ilvana Dzafic, Roshini Randeniya, Clare D. Harris, Moritz Bammel, Marta I. Garrido

**Affiliations:** Melbourne School of Psychological Sciences, University of Melbourne, Parkville, VIC, Australia; Queensland Brain Institute, University of Queensland, Brisbane 4072, Australia; Australian Research Council Centre of Excellence for Integrative Brain Function, Australia; Centre for Advanced Imaging, University of Queensland, Brisbane 4072, Australia; Institute of Cognitive Science, University of Osnabrueck, Osnabrueck, Germany

**Keywords:** prediction error, volatility, psychosis continuum, sensory learning, inference

## Abstract

Our perceptions result from the brain’s ability to make inferences, or predictive models, of sensory information. Recently, it has been proposed that psychotic traits may be linked to impaired predictive processes. Here, we examine the brain dynamics underlying sensory learning and inference in stable and volatile environments, in a population of healthy individuals (N=75) with a range of psychotic-like experiences. We measured prediction error responses to sound sequences with electroencephalography, gauged sensory inference explicitly by behaviourally recording sensory ‘regularity’ learning errors, and used dynamic causal modelling to tap into the underlying neural circuitry. We discuss the findings that were robust to replication across the two experiments (N=31 and N=44 for the discovery and the validation datasets, respectively). First, we found that during stable conditions, participants demonstrated a stronger predictive model, reflected in a larger prediction error response to unexpected sounds, and decreased regularity learning errors. Moreover, individuals with attenuated prediction errors in stable conditions were found to make greater incorrect predictions about sensory information. Critically, we show that greater errors in sensory learning and inference are related to increased psychotic-like experiences. These findings link neurophysiology to behaviour during sensory learning and prediction formation, as well as providing further evidence for the idea of a continuum of psychosis in the healthy, non-clinical population.

**Significance Statement:** Whilst perceiving the world, we make inferences by learning the *regularities* present in the sensory environment. It has been argued that psychosis may emerge due to a failure to learn sensory regularities, resulting in an impaired representation of the world. Recently it has been proposed that psychosis exists on a *continuum*; however, there is conflicting evidence on whether sensory learning deficits align on the *non-clinical* end of the psychosis continuum. We found that sensory learning is associated with brain prediction errors, and critically, it is impaired in healthy people who report more psychotic-like experiences. We replicated these findings in an independent sample, demonstrating strengthened credibility to support that the continuum of psychosis extends into the non-clinical population.

## Introduction

In a stable environment, sensory perception is facilitated by prior beliefs about what is likely to happen next ^1, 2^. By estimating the probability of events given the past history (i.e. inference based on a learnt regularity), we can form a predictive model about what is likely to happen next^3^. When circumstances are ‘volatile’, such that previously learnt regularities change erratically, it is advantageous to form more flexible predictive models ^4, 5^. Indeed, previous literature has shown that healthy individuals are able to estimate environmental volatility ^5^, adopting a greater learning rate in the face of ever changing, volatile circumstances ^6^. This motivates exploratory behaviour and continuous updating, as well as suppressing of top-down prior beliefs ^7^. However, the underlying brain dynamics involved during sensory learning in volatile environments, and that support volatility attuning, are currently not known.

The state of constant learning is inefficient as a long-term strategy in environments that are stable ^8, 9^. Stable environments allow for the development of a robust predictive model, which simultaneously enables an efficient encoding of sensory stimuli while minimizing the demand on cognitive resources^10^. In these environments, healthy individuals form strong predictions about forthcoming sensory stimuli, and their brains consequently produce large prediction error (PE) responses to events that violate such predictions^11-13^. The PE response is commonly gauged using electroencephalography (EEG) and an auditory oddball paradigm ^14^, in which surprising sounds are embedded in a sequence of predictable sounds. PE responses are thought to signify *implicit* regularity learning ability: an individual’s accuracy in their inherent learning of the statistics of sensory events ^15, 16^. However, it has not as yet been addressed if PE responses scale with the accuracy in sensory learning and inference.

Emerging theoretical accounts of psychosis postulate that psychotic experiences arise due to an impairment in the brain’s predictive ability to infer internal and external sensations ^17-19^. Converging lines of evidence support the theory that psychosis arises due to an impaired predictive model of sensory events ^20, 21^. The most robust and replicable empirical evidence for this arises from attenuated PE responses in psychosis ^15, 17, 22, 23^. Alterations in the neurophysiology of PEs have been shown to increase as psychotic traits increase, suggesting that the degree of PE aberrancy aligns on a continuum of psychosis ^24, 25^. However, the underlying brain networks, presumably also altered along the continuum, remain unknown. The psychosis continuum comprises the full spectrum of psychotic experiences, from healthy individuals who experience a range of psychotic-like experiences, to prodromal individuals with subclinical symptoms, and to those with florid psychosis at the very end of the spectrum ^26, 27^. In this study, we zoom in to the non-clinical population, as there is conflicting evidence whether PEs are also reduced on the healthy end of the continuum ^28, 29^. One of the benefits of investigating psychotic-like experiences ^30, 31^ in the non-clinical end of the spectrum is the possibility to eschew the confounds of medication and illness severity.

The aims of this study were three-fold. Firstly, we wanted to elucidate the neural underpinnings of sensory learning in different volatility contexts. For this purpose, we developed a novel auditory oddball task with either fixed sound probabilities (stable conditions) or varying sound probabilities (volatile conditions; see Figure 2). The second aim was to examine the relationship between regularity learning behaviour, sensory PEs and the brain’s ability to attune to volatility. The third aim was to investigate whether aberrations in sensory learning and the underlying effective connectivity are aligned on the non-clinical continuum of psychosis. In our discovery study we found significant findings supportive of all our hypotheses described above. However, in response to the current replication crisis in the field, we sought to corroborate these findings in an independent sample. In so doing, we hoped to assess the robustness of our initial findings, using the same methodology in an unbiased manner. To this end, we report our findings side-by-side, for the discovery and the validation samples, and discuss the implications of findings that were replicated.

**Fig. 1.**
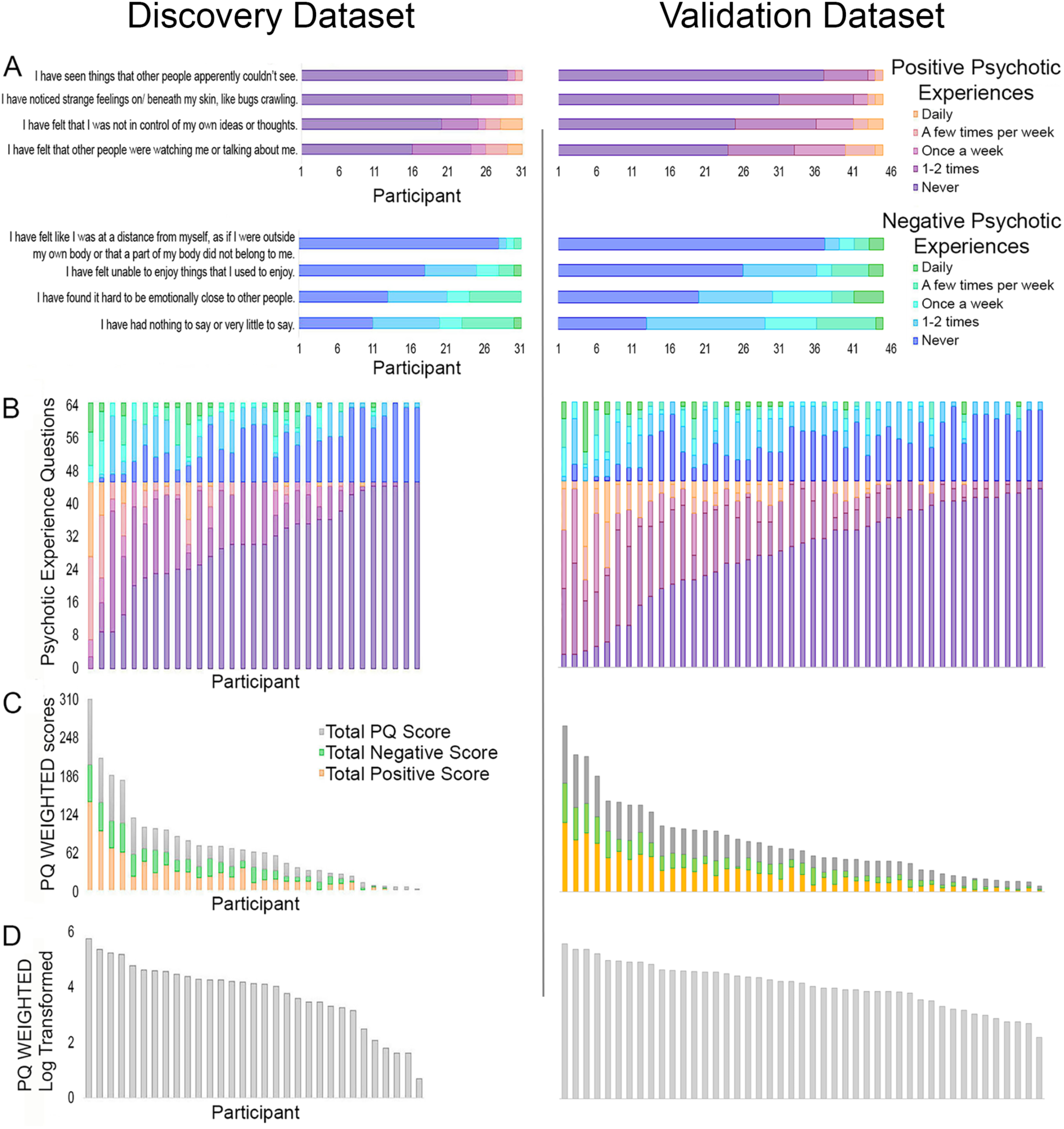
Prodromal questionnaire (PQ) scores for discovery and validation samples. A) Frequency of a subset of positive psychotic experiences (purple to orange) and negative psychotic experiences (blue to green), B) Frequency of psychotic experiences, displaying positive psychotic experiences (purple to orange) and negative psychotic experiences (blue to green), C) PQ scores weighted by frequency, displaying total PQ weighted scores (grey), negative weighted scores (green) and positive weighted scores (orange), D) Log transformed PQ weighted scores.

**Fig. 2.**
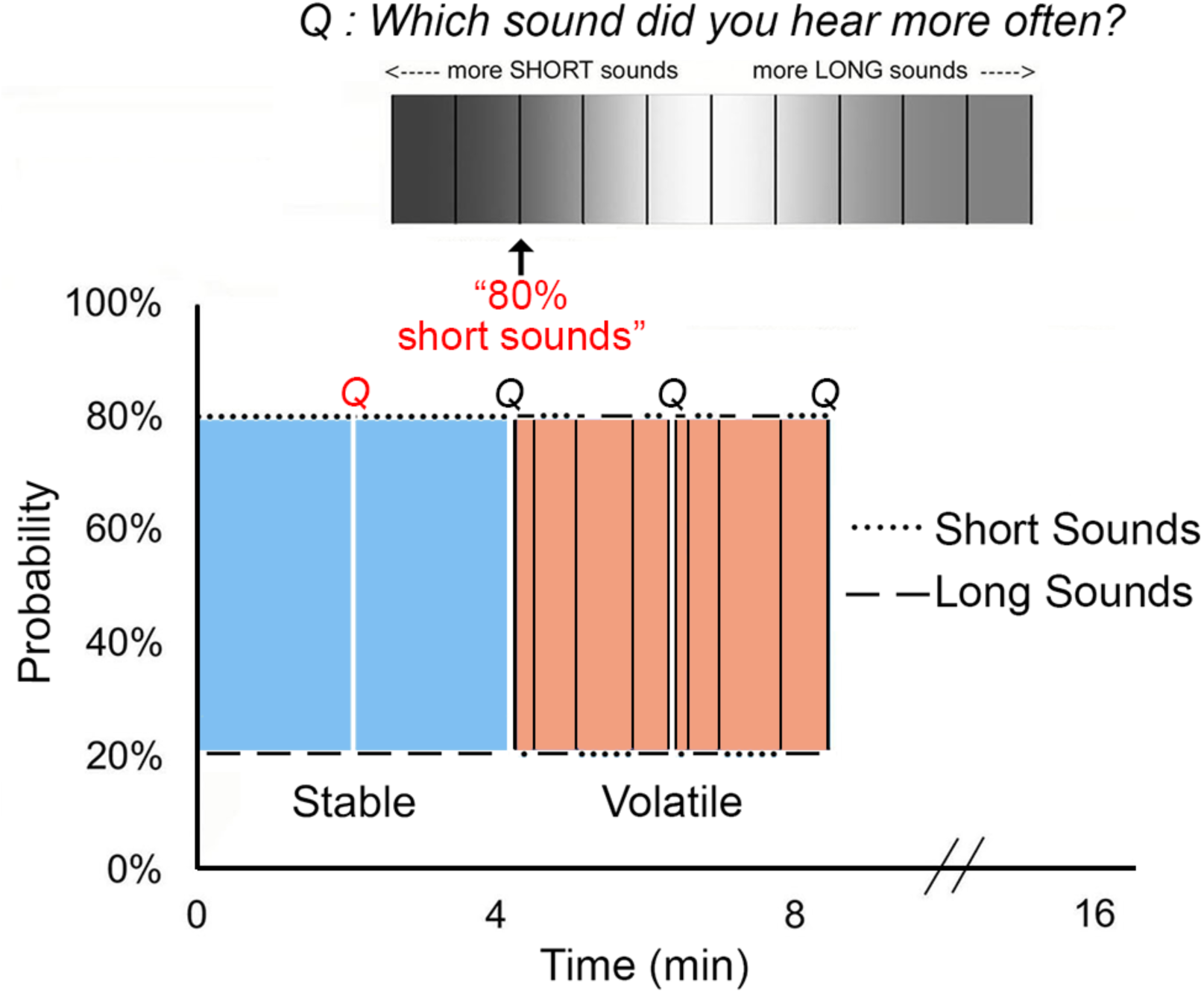
Reversal oddball task. A schematic diagram showing an example of a stable and volatile run. In the stable run depicted here, the short sound (50 ms) was more probable (80%) throughout, whereas in the volatile run the long sound (100 ms) was more probable in the first block, then the short sound was more probable in the second block, with eight blocks and seven reversals of probability in total. Participants were asked to listen to the sounds and estimate the probability (top Q) of the most frequent sound and rate their confidence on this judgment, every 2:08 min.

## Methods and Materials

### Participants

The total sample from both the discovery study and the replication included seventy-five healthy adults. The discovery study included thirty-one participants (age range: 19 to 38 years; mean age: 24.65 years, *SD* = 4.85; 14 males and 17 females) and the replication study included forty-four participants (age range: 18 to 39 years; mean age: 24.27 years, *SD* = 5.13; 22 males and 22 females). All participants were recruited through the Psychology Research Participation Scheme (SONA), an online newsletter to staff and students across the University of Queensland and Gumtree. Prior screening confirmed that all participants did not have a history of psychiatric or neurological disorders, and were not currently taking medication or using any illicit drugs. The highest level of education, smoking habits and alcohol consumption were recorded. Participants also completed the 92 Item - Prodromal questionnaire (PQ), which measures positive and negative symptoms and is typically used to assess psychotic experiences in healthy individuals ^32^. For further information on the demographics of the sample and the PQ scores per participant, as well as the positive and negative symptom frequencies, please see Figure 1, and in the supplementary materials Table S1. Participants provided written informed consent for taking part in our study after reading and understanding the information sheet, which included a full description of the study and procedure. Participants received monetary reimbursement for their time. Participant recruitment and data collection for the discovery and validation samples were conducted by independent researchers in the same lab using the same methodology. This research was approved by the University of Queensland Human Research Ethics.

### Materials and Procedure

#### Reversal oddball task

An auditory ‘duration’ oddball paradigm was modified so that the probability of different sounds varying in duration was either stable or volatile (adapted from Weber and colleagues^33^). In a stable experimental run, a particular sound was always more likely than another sound (e.g. short sounds had 80% probability and long sounds had 20% probability). In volatile experimental runs, a particular sound, which was more likely in the first block, was then less likely in the second block, with four blocks and three reversals of probability in total (the stable experimental runs only included one block). The Reversal oddball task is represented in Figure 2.

#### Auditory stimuli and experimental design

The Reversal oddball paradigm consisted of 2000 pure tones played over eight experimental runs (2:08 min each; four stable and four volatile runs). The tones varied in duration such that short tones lasted 50 ms and long tones lasted 100 ms. All tones had an identical frequency of 500 Hz and a smooth rise and fall periods of 5ms. The tones were presented in a pseudorandom order, with each presentation of 5 tones including a deviant tone in a randomly assigned position; the deviant tones were always separated by at least one standard tone. The tones were delivered binaurally via insert earphones for ∼2 min every 500 ms. Sound intensity was kept constant between participants at a comfortable level. The order of stable and volatile runs was counterbalanced across participants.

#### Procedure

During the Reversal oddball task participants were seated on a comfortable chair in front of a desk and computer screen, in a dimly lit Faraday cage testing room. Prior to the experiment, the participants were familiarized with the different sound types and trained with two short practice runs of the task. Participants were asked not to move while the sounds were played and to look at a fixation cross at the centre of the screen. The participants were instructed to pay attention to the sounds in order to judge the proportion of different sound types and rate their confidence on this judgment. Participants were required to make these estimates every 2:08 min using a computer keyboard and a mouse. The total duration of the Reversal oddball task was approximately 20 minutes (including short breaks).

### EEG Recording and Pre-processing

A Biosemi Active Two system recorded continuous electroencephalography (EEG) data from 64 scalp electrodes at a sampling rate of 1024Hz. Electrodes were arranged according to the international 10-20 system for electrode placement (Oostenveld, R. & Praamstra, P, 2001). Standard pre-processing and data analysis were performed with SPM12 (http://www.fil.ion.ucl.ac.uk/spm/). Full details regarding the pre-processing steps can be found in the supplementary materials.

### Data Analysis

In the current study we employed both frequentist and Bayesian approaches to our analyses. A brief explanation of the Bayesian approach can be found in the supplementary materials. The Bayesian analyses were conducted using the JASP package (https://jasp-stats.org/). The frequentist analyses were conducted using the SPSS package (IBM Corp, 2012); multiple correlations were corrected using the Šidák method ^34^. In the validation study we excluded 2 participants in the neuroimaging analyses due to EEG trigger failure and high impendence (> ±50 Ω). Further, 1 participant was removed from the ERP analyses, and 1 participant from the PQ analyses, as they were an outlier (z-score > ±3).

We conducted single-channel event-related potential (ERP), whole-channel spatiotemporal, source level, and effective connectivity analyses in order to assess the effect of environment (Stable vs. Volatile) on neuronal activity. In addition, we conducted behavioural analyses to investigate regularity learning in the different volatility contexts, and its associations with psychotic-like experiences and neural dynamics. Full details regarding the single-channel and behavioural analyses, as well as the spatiotemporal, source reconstruction, and dynamic causal modelling analysis can be found in the supplementary materials. Finally, we conducted Pearson’s and Bayesian correlations to assess the association between psychotic-like experiences and the effective connectivity estimates. In the discovery study, the top-down, right frontotemporal (right IFG to right STG) connection demonstrated a strong, significant association with psychotic experiences (*p* = 0.005, BF_-0_ = 18.28; see Table S3). The association was followed up with a mediation analysis using Preacher and Hayes ^35^ bootstrap method (PROCESS in SPSS) to directly investigate whether top-down connectivity was a mechanism by which regularity learning influenced the severity of psychotic-like experiences ^36^.

## Results

Our first aim was to compare the strength of the predictive models established in stable and volatile contexts in a regularity learning task. For this purpose we examined the event-related potential (ERP) recorded at frontocentral channel (Fz). In line with the vast oddball literature we consistently found that responses to deviant sounds were larger than responses elicited by standard sounds, regardless of volatility for both the Discovery (*F*(1,30) = 45.33, *p* < 0.001, η^2^ = 0.60) and the Validation datasets (*F*(1,40) = 88.14, *p* < 0.001, η^2^ = 0.69). Moreover, we found a significant interaction between PE response and volatility in the Discovery dataset (*F*(1,30) = 11.06, *p* = 0.002, η^2^ = 0.27), which was replicated in the Validation dataset (*F*(1,40) = 12.26, *p* = 0.001, η^2^ = 0.24). Critically, a follow-up analysis revealed that PEs were larger under the stable compared to the volatile conditions again for both the Discovery (*t*(30)=-3.33, *p* = 0.002, *d* = −0.60) and the Validation sets (*t*(40)=-3.74, *p* = 0.001, *d* = −0.61; see Figure 3a and 3b).

**Fig. 3.**
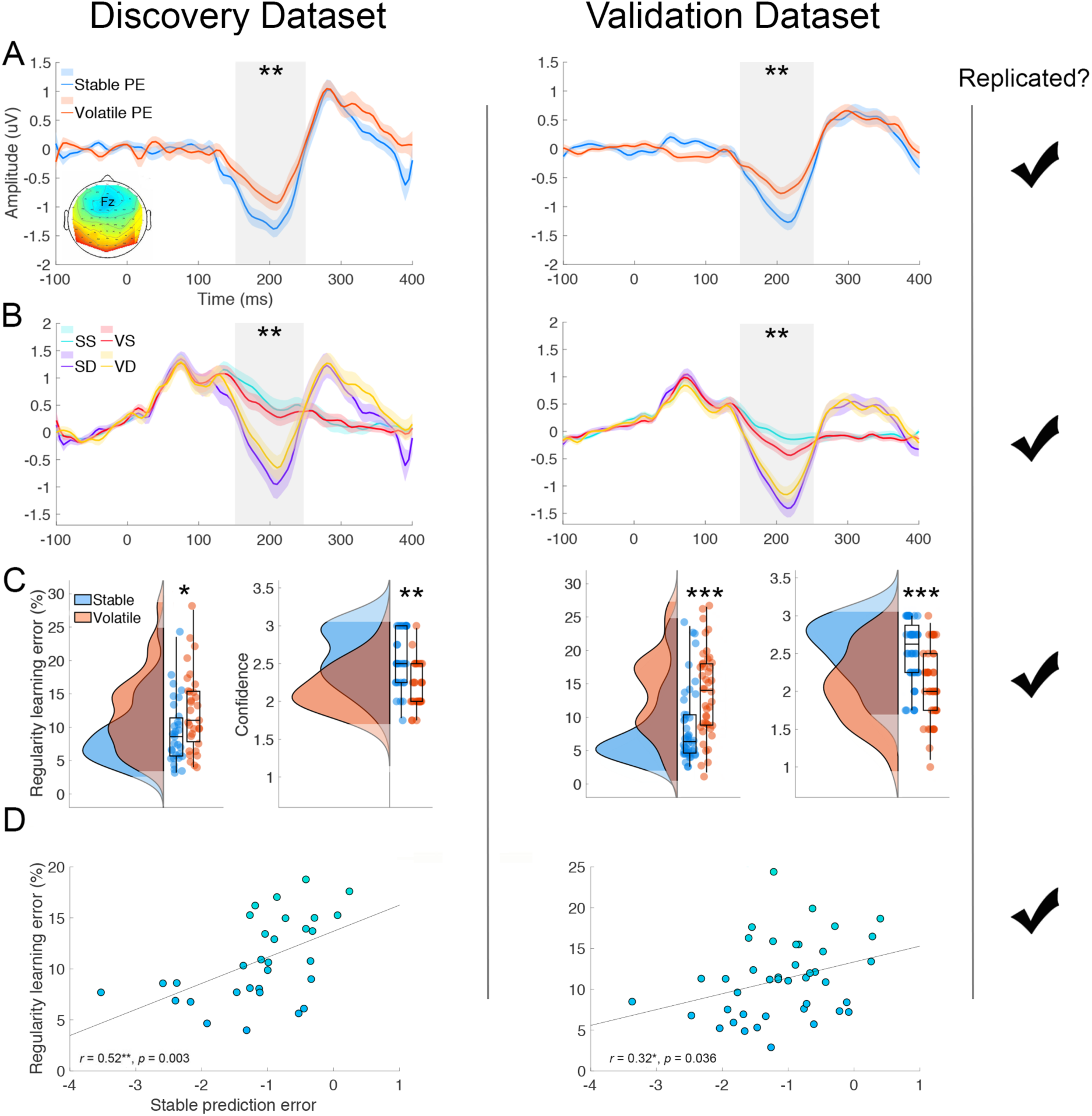
Prediction errors relate to regularity learning ability indicating a stronger prediction model in *stable* conditions. A) Significant interaction between volatility (stable – blue, versus volatile – orange) and brain PE responses, B) Significant main effect of PE, showing brain responses evoked by standards (green and red) and deviants (purple and yellow) in the context of stable and volatile conditions, C) Fewer regularity learning errors and greater confidence in stable (blue) than volatile (orange) conditions, D) Significant correlation between regularity learning errors and PEs in stable conditions. SS = Stable standard; SD = Stable deviant; VS = Volatile standard; VD = Volatile deviant. * p < 0.05; ** p < 0.001, *** p < 0.0001. All of these findings were replicated across the discovery and the validation datasets.

We next examined differences in mean percentage errors in probability estimation (regularity learning error) and mean confidence ratings during stable and volatile conditions. The data showed that participants had fewer errors in regularity learning during stable conditions (Discovery: *M* = 9.30%, *SE* = 0.87; Validation: *M* = 8.62%, *SE* = 0.87), compared to volatile conditions (Discovery: *M* = 12.38%, *SE* = 1.07; Validation: *M* = 13.54%, *SE* = 0.97), Discovery: *t*(30) = −2.40, *p* = 0.023, *d* = −0.43; Validation: *t*(43) = −4.21, *p* < 0.0001, *d* = −0.81. In addition, participants had greater confidence in their probability estimates during stable conditions (Discovery: *M* = 2.57, *SE* = 0.07; Validation: *M* = 2.53, *SE* = 0.06), compared to volatile conditions (Discovery: *M* = 2.19, *SE* = 0.05; Validation: *M* = 2.08, *SE* = 0.07), Discovery: *t*(30) = 4.78, *p* < 0.001, *d* = 0.86; Validation: *t*(43) = 6.98, *p* < 0.0001, *d* = 1.00 (see Figure 3c).

We then asked whether regularity learning errors were related to the degree of PE response. Pearson’s correlations and Bayesian analysis revealed a significant correlation between regularity learning errors and PEs (at the ERP level) in stable conditions (Discovery: *p* = 0.003 (*p*_adjusted_ < 0.01), BF_+0_ = 33.99; Validation: *p* = 0.036, BF_+0_ = 3.14; see Figure 3d, Table S2), which had very strong evidence in the discovery dataset and moderate evidence in the validation dataset.

To further investigate the sensory PEs evoked by regularity violations with fewer spatial and temporal constraints, we ran a general linear model for the whole spatiotemporal volume of brain activity. Firstly, we supported previous auditory oddball findings by showing a significant main effect of PE response (standard sounds vs deviant sounds) peaking at 290 ms (peak-level *F* = 240.99, *p*_FWE_ < 0.001) and 205 ms (peak-level *F* = 170.92, *p*_FWE_ < 0.001) in central and occipitoparietal channels, and 25 ms (right frontal channels; peak-level *F* = 25.26, *p*_FWE_ = 0.004) in frontal channels. We also replicated our own findings by showing main effects of PE response in the validation dataset that peaked at similar time points: 300 ms (peak-level *F* = 176.59, *p*_FWE_ < 0.001), 210 ms (peak-level *F* = 77.44, *p*_FWE_ < 0.001) and 185 ms (peak-level *F* = 68.76, *p*_FWE_ < 0.001) in central and occipitoparietal channels (note: 25 ms was not replicated). We originally found a significant interaction between PE response and volatility, at 165 ms over occipitocentral channels (peak-level *z* = 4.27, *p*_FWE_ = 0.015, see Figure 4a); however, this did not replicate in the validation set.

**Fig. 4.**
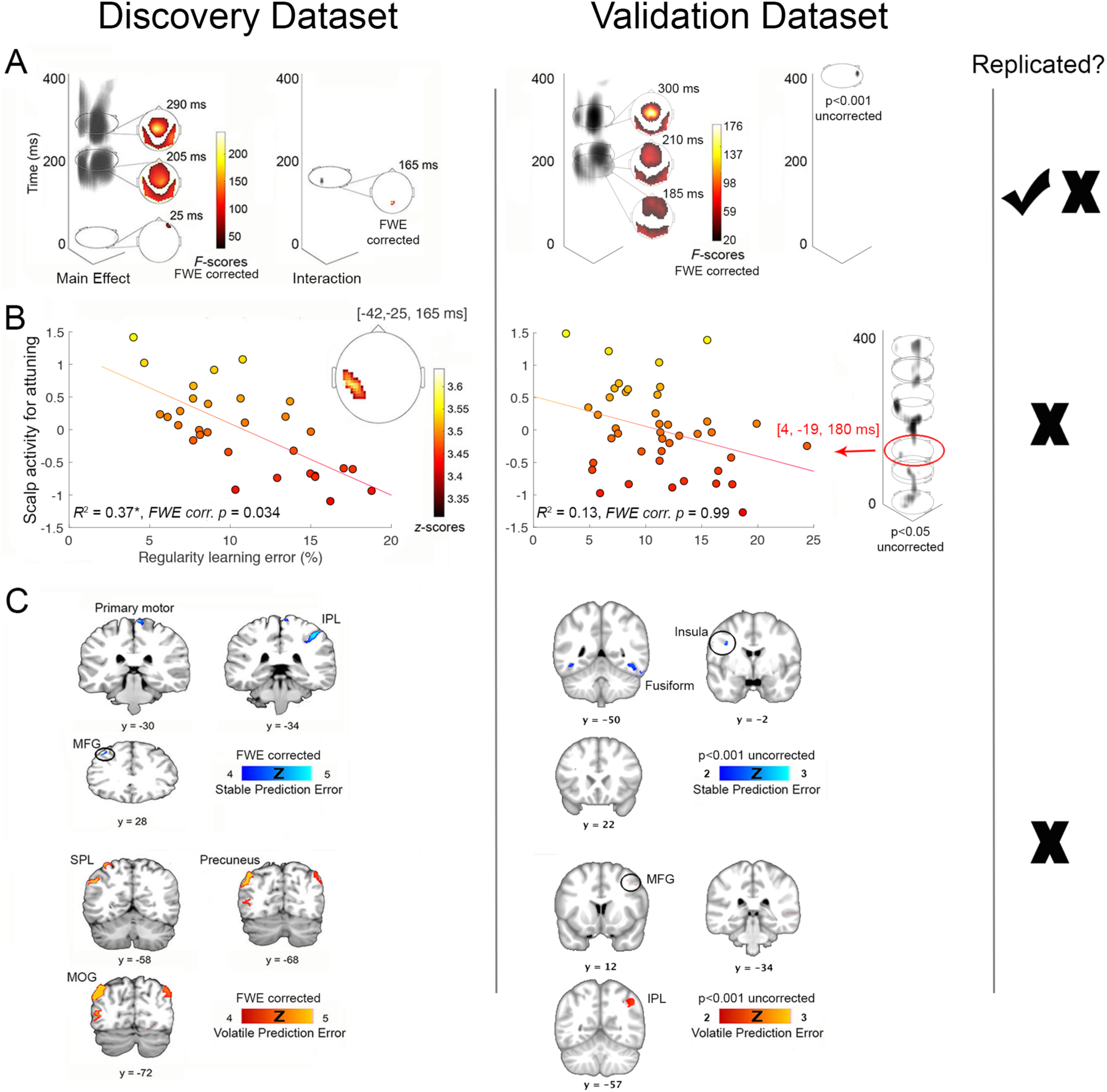
Brain responses underlying main effect of prediction error. A) Spatiotemporal univariate analysis revealed a significant main effect of PE (left column; this was replicated); and PE x volatility interaction (right column; this was not replicated); B) Spatiotemporal multiple regression analysis revealed a negative relationship between regularity learning errors and activity during the interaction (Stable PE > Volatile PE); however, this result was not replicated at FWE corrected significance, even after a region of interest analysis; C) Source reconstruction analysis revealed significant clusters for stable PEs versus volatile PEs; however, these results were not replicated. IPL = inferior parietal lobule; MFG = middle frontal gyrus; SPL = superior parietal lobule; MOG = middle occipital gyrus. Except where otherwise specified, all maps are displayed at *p* < 0.05, FWE whole-volume corrected.

Next, we asked whether regularity learning errors were related to the neuronal correlates of volatility attuning. To address this question, we conducted a spatiotemporal multiple regression analysis of the interaction between PEs and volatility (Stable PEs > Volatile PEs) with regularity learning error as the predictor variable. Our data in the discovery study showed that a decrease in regularity learning errors significantly predicted an increase in brain activity at 165 ms (peak-level *z* = 3.64, cluster-level *p*_FWE_ = 0.034, see Figure 4b). However, this result did not replicate, even after conducting a spatiotemporal volume of interest analysis (based on the result from the discovery study) for the multiple regression.

In the discovery study, source-level analysis revealed that stable PEs engaged the middle frontal gyrus (peak-level *z* = 4.23, *p*_FWE_ = 0.02), primary motor area (peak-level *z* = 4.45, *p*_FWE_ = 0.009), and inferior parietal lobule (peak-level *z* = 4.56, *p*_FWE_ = 0.006). In comparison, volatile PEs engaged precuneus (peak-level *z* = 5.05, *p*_FWE_ = 0.001) and middle occipital gyrus (peak-level *z* = 4.39, *p*_FWE_ = 0.011). However, we were not able to replicate these findings in the independent validation sample. None of the voxels survived FWE correction, nor did the uncorrected clusters correspond with the discovery study (see Figure 4c).

The network architecture underlying PE response has been extensively studied previously ^37, 38^. Here, we were interested in, 1) the effect of contextual volatility on neuronal PE responses, and 2) the effective connectivity underpinning psychotic traits and sensory learning.

Bayesian model comparison was performed on thirty-six different dynamic causal models (see Figure S3), which were based on the functional brain architecture previously shown to underlie PE responses ^37, 38^. Here, PEs in stable blocks were compared to volatile blocks. Results from Bayesian model selection using random effects family-level analysis (in both the discovery and validation datasets) indicated that the best model included connections amongst six *a priori* defined regions, with inputs to left and right primary auditory cortex (A1), intrinsic connections within the A1, bilateral connections between A1 and superior temporal gyri (STG) and between STG and inferior frontal gyri (IFG), as well as lateral connections between left and right A1, and left and right STG (model 7; see Figure 5a). The optimal model, which had greater modulation in backward, forward and intrinsic connections, was replicated across both the discovery and the validation datasets (see Figure 5a).

**Fig. 5.**
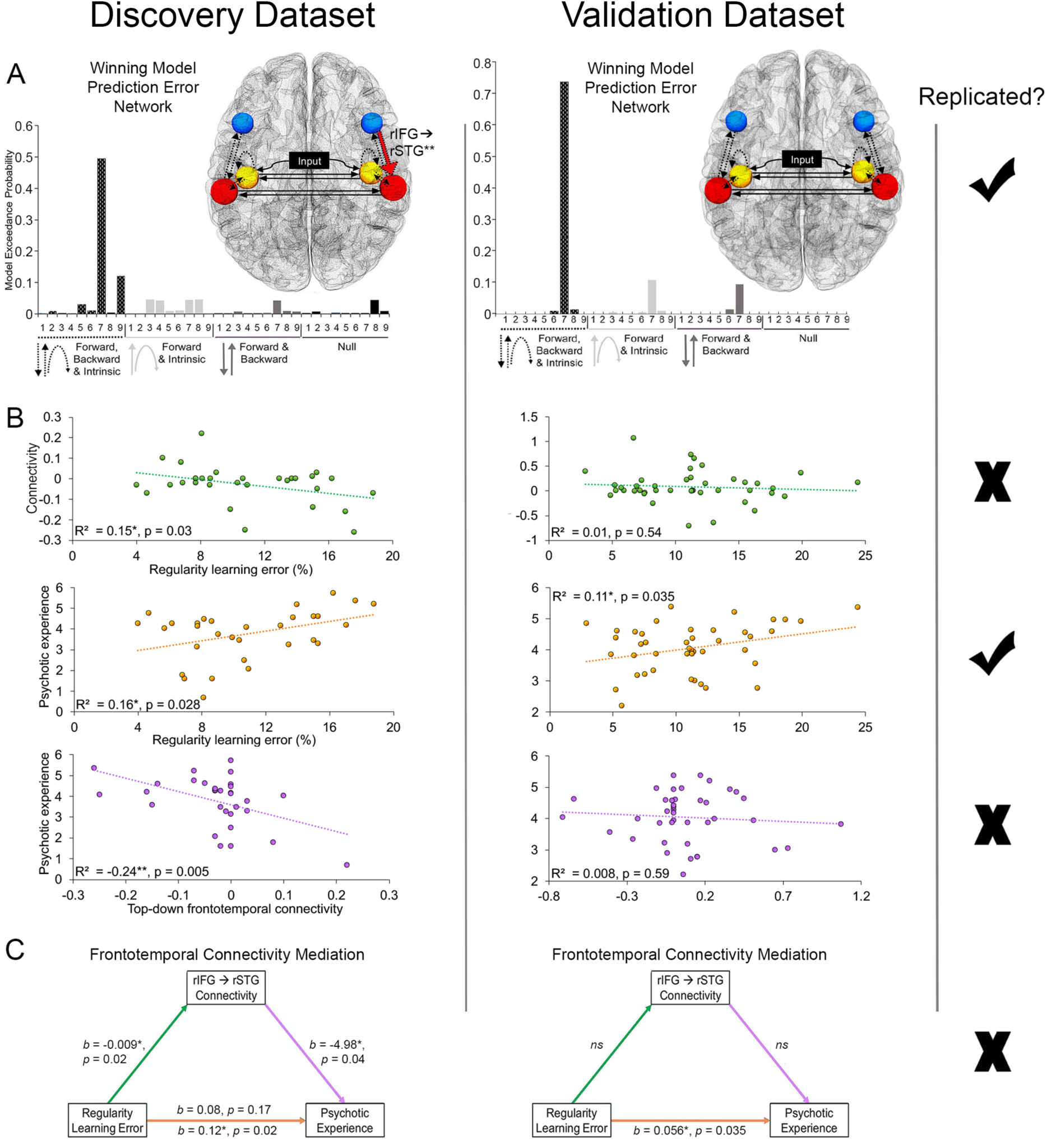
Regularity learning errors predict severity of psychotic-like experiences in the non-clinical population. A) The winning model architecture, replicated across both datasets, had connections between all six regions bilaterally (model 7), included volatility-dependent modulations in forward, backward and intrinsic connections; B) Regression plots for regularity learning errors and top-down frontotemporal connectivity (in green; not replicated); regularity learning errors and psychotic-like experiences (in orange; replicated); and top-down frontotemporal connectivity and psychotic experiences (in purple; not replicated); C) The mediation analysis revealed a significant full mediation in the discovery sample, with an indirect effect of regularity learning on psychotic experience through top-down frontotemporal connectivity; however, this finding was not replicated in the validation sample.

In order to test the evidence for a continuum of psychosis, we examined the altered neural dynamics, behaviour and neurophysiology related to psychotic traits in the general non-clinical population. First, we examined brain connectivity estimates by applying Bayesian model averaging across all models (weighted by their probability) and participants. In the discovery dataset, we found a strong, significant negative correlation between psychotic experiences and top-down connectivity from the right IFG to STG (frontotemporal) (*p* = 0.005 (*p*_adjusted_ < 0.05), BF_-0_ = 18.28; see Table S3). However, this was not replicated. Next, we asked if aberrations in behaviour (greater regularity learning errors) and neurophysiology (attenuated PE) were also aligned on the psychosis continuum. Pearson’s and Bayesian correlations were conducted on psychotic experiences, regularity learning errors, as well as PEs (at the ERP level) in stable and volatile conditions. We found a significant correlation between psychotic experience and errors in regularity learning that was replicated across both datasets (Discovery: *p* = 0.028, BF_+0_ = 4.37; Validation: *p* = 0.035, BF_+0_ = 2.55; see Table S2). This showed that healthy individuals with greater psychotic experiences were worse at learning about sensory regularities. The fact this was replicated provides strong support for behavioural alterations in sensory inference aligning on the non-clinical end of the psychosis continuum.

Our final analysis explored if the top-down frontotemporal connection, which was weaker in individuals with more psychotic experiences in the discovery study, mediated the relationship between regularity learning errors and psychotic traits. For this purpose, we employed a mediation analysis ^36^. Multiple regressions were conducted to asses each component of the mediation analysis (see Figure 5c). In the discovery dataset, we found that regularity learning error was a significant predictor of top-down frontotemporal connectivity (*b* = −0.009, *p* = 0.02), and that top-down frontotemporal connectivity was a significant predictor of psychotic experience (*b* = −4.98, *p* = 0.04). We found a significant indirect effect (*ab* = total effect - direct effect) of regularity learning on psychotic experience through top-down frontotemporal connectivity (*ab* = 0.05, Bias Corrected and Accelerated Bootstrap (BCA) CI [0.004, 0.14], P_M_ = 0.38 - percent mediation: percent of the total effect accounted for by the indirect effect). Regularity learning was no longer a significant predictor of psychotic experiences after controlling for the mediator: top-down frontotemporal connectivity (*b* = 0.08, *p* = 0.17), consistent with a full mediation (see Figure 5b). However, the mediation hypothesis was only supported (and conducted) in the discovery dataset as we failed to replicate the previously observed significant relationships between top-down connectivity and regularity learning errors as well as between top-down connectivity and psychotic experiences.

## Discussion

The aims of the current study were to investigate and replicate the neural mechanisms that underpin sensory learning and volatility attuning in healthy individuals with a range of psychotic-like experiences. We measured individuals’ regularity learning abilities (by asking them to estimate the probabilities of sounds) in stable and volatile conditions while recording their brain activity using EEG. We pursued these goals in two independent datasets and in turn discuss the findings that did replicate across both. During stable conditions, compared to volatile, regularity learning improved, prediction errors increased, and there was greater modulation of intrinsic, forward and backward connections. For the first time, we showed that regularity learning (behaviourally assessed) relates to prediction errors. Moreover, we were able to replicate the finding that a greater degree of psychotic-like experiences in healthy individuals is associated with impaired sensory regularity learning ability, providing strong evidence for the existence of a *continuum of psychosis* in the non-clinical population.

PE responses, regularity learning and confidence were enhanced in *stable*, more predictable environments, than in more volatile, less predictable, environments. Increased PEs in stable (compared to volatile) conditions have been identified in previous studies ^39, 40^. PEs signal a violation between what the brain predicts will happen and what is actually experienced. As such, PEs are fundamental teaching signals that drive updating of the brain’s predictive model of the sensed world ^41, 42^. Thus, greater PE responses to regularity violations indicate a stronger (i.e., more precise) prediction model in stable than volatile conditions ^42, 43^. Indeed, PEs in stable conditions were found to be associated with better sensory regularity learning in the current study. Regularity learning is the process by which the brain learns the statistical structure in the environment and forms predictive models of what is likely to happen next ^44-46^. Previous studies have demonstrated that individuals are able to implicitly learn the statistical structure of sensory events in the environment ^3, 15^. Crucially, by simultaneously recording PE responses and behaviourally measuring regularity learning, we showed, for the first time and across both the discovery and the validation samples, that greater sensory PEs are associated with improved explicit ability to gauge the sensory regularities within one’s environment. This was specifically found for PEs in *stable* rather than volatile conditions, suggesting that PE’s (i.e. MMN) may not relate to the long-term rule updates (i.e. global regularities) that influence regularity learning in the volatile conditions.

At the neural level, we found differences in the pattern of connections underlying PE response in stable compared to volatile conditions. Larger PEs in stable compared to volatile conditions were produced by enhanced modulations in backward, forward and intrinsic connections. Forward and backward connections are thought to convey PEs and predictions (i.e., beliefs about sensory input), respectively. Intrinsic connections emulate local adaptation of neuronal responses and are thought to reflect the precision (strength) of neuronal representations ^47^. This finding is in keeping with the predictive coding account of the mechanisms underlying perception of an auditory oddball sequence ^48^, and suggest that more precise predictive models about sensory input are enabled by greater brain connectivity in stable than volatile PEs.

We did not find consistent associations between alterations in the neurophysiology or brain connectivity and psychotic-like experiences in our non-clinical samples. While, in our discovery sample we found weaker top-down frontotemporal connectivity in people that reported more psychotic-like experiences, we failed to replicate this finding in the validation sample. It is possible that these brain alterations are not as robust in the healthy population that experience mild, psychotic-like experiences. Indeed, there is conflicting evidence regarding attenuation of PEs (such as MMN and P300) in the non-clinical psychosis continuum. A recent study has found reductions in sensory PEs ^29^; while others have reported a lack of/mixed evidence for an association ^28, 49, 50^. Further research looking at the full continuum including a range of psychiatric groups may elucidate whether or not a relationship between psychotic experiences and brain alterations does exists; and if so, whether it also extends into the non-clinical, healthy population.

Critically, in our study we uncovered the *behavioural* aberrations underlying predictive processing that are aligned on the non-clinical continuum of psychosis. Previous research investigating behaviours such as force-matching, associative learning and reversal learning have failed to replicate the relationship between altered behaviour and psychotic-like experiences in the general population ^51^. In response to the recent replication crisis in the field, we decided to collect two independent samples for discovery and validation of our findings. Indeed, we were able to replicate that psychotic-like experiences are associated with impaired regularity learning (measured behaviourally). Therefore, we show, with strengthened credibility, that there exists a continuum of psychosis, even in the non-clinical end of the spectrum, that manifests on a declining ability in regularity learning.

### Conclusion

In the current study, we explored the brain dynamics underpinning regularity learning under uncertainty, as well as the relationship between disruptions to predictive processes and psychotic-like experiences in healthy individuals. We originally investigated these processes in a discovery dataset and then sought to replicate the findings in an independent validation dataset. We found that individuals learn better and their brain PE responses are greater during stable than volatile conditions. There is greater modulation of intrinsic, forward and backward connections in stable conditions, when a more precise model of the sensed environment is represented in the brain. In addition, we were able to show that individuals with stronger stable PE responses had improved sensory regularity learning behaviour. Importantly, we were able to replicate that aberrations in predictive processes are aligned on a non-clinical continuum of psychosis, in the sense that healthy people with more psychotic-like experiences have poorer regularity learning ability.

## Acknowledgements

The authors thank the participants for their time.

This work was funded by the Australian Research Council Centre of Excellence for Integrative Brain Function (ARC Centre Grant CE140100007) and a University of Queensland Fellowship (2016000071) to MIG, as well as a University of Queensland International Research Scholarship to RR.

## Data and materials availability

The article’s supporting data and materials have been made available. Please find raw data files and the behavioural/connectivity scores here: https://espace.library.uq.edu.au/view/UQ:724759.

