## Supplementary for "Sensory learning and inference is impaired in the non-clinical continuum of psychosis: a replication study"

**This PDF file includes:**

Supplementary text

Figs. S1 to S3

Tables S1 to S3

References for SI reference citations

**Supplementary Information Text**

**EEG preprocessing.** Data were referenced to standard BioSemi reference electrodes, down-sampled to 200Hz and high-pass filtered at 0.5 Hz using the Butterworth filter. Eye blinks were detected and marked using the VEOG channel at an eyeblink threshold of 4; the Berg method was used to correct for eye blinks. The data were epoched offline with a peri-stimulus window of -100 to 400ms. Further artefact rejection was performed by thresholding all channels at 100uV, robustly averaging across trials (1), applying a low-pass Butterworth filter of 40 Hz, and baseline correcting between -100 to 0 ms. We analysed event-related potentials from the onset of standard and oddball tones, separately for stable and volatile conditions.

**Bayesian approach.** Bayes factors are based on Bayes’ rule, displayed in the equation below. The posterior probability given the observed data (the posterior; p(H_1_|data)/p(H_0_|data)), equals the prior odds (the odds of the null and alternative hypotheses (p(H_1_)/p(H_0_)) before the data are observed), multiplied by the Bayes factor (p(data|H_1_)/p(data|H_0_)), or the change (update) from prior to the posterior (2).

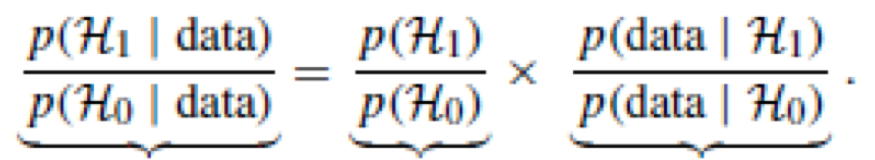

The Posterior	The Prior	Bayes factor BF_10_

The subscript ‘10’ in BF_10_ indicates that in the equation H_1_ (the alternative hypothesis) is in the numerator and H_0_ (the null hypothesis) is in the denominator, and subscript ‘01’ indicates the reverse. In the current study, Bayesian factors were computed using BF_10_, which indicates testing the alternative hypothesis over null hypothesis.

**Single-channel and behavioral analyses.** We conducted a full factorial 2x2 within subjects ANOVA design on mean ERP values with Environment (Stable and Volatile) and PE (Standard and Deviant) as factors. Mean ERP values were obtained by averaging across the preselected time window of interest, which is typical for PE latency: 150 – 250 ms, over a frontocentral channel (Fz), in which PE responses are typically seen in oddball paradigms (3). We contrasted evoked responses to deviant and standard sounds, under stable and volatile conditions. Significant interactions were further analyzed using paired t-tests.

We conducted paired t-tests on mean percentage errors in probability estimation (a proxy for regularity learning) and mean confidence in probability estimation, in stable vs. volatile conditions. This was done in order to assess the effect of environment (Stable vs. Volatile) on regularity learning and confidence in estimating probabilities. Next, we computed Pearson’s and Bayesian correlations to assess the association between psychotic experience, regularity learning errors, and PEs in stable and volatile conditions (see Table S2).

**Spatiotemporal maps and source reconstruction.** Three-dimensional spatiotemporal images were generated from averaged ERP data for each participant and condition. A two-dimensional matrix, corresponding to the scalp electrode space was produced, for each time bin from 0 to 400ms in steps of 5ms. The images were assembled according to their peristimulus temporal order, which resulted in a three-dimensional spatiotemporal image (32 × 32 × 81) per participant. These images were then smoothed at full width half maximum of 12 mm × 12 mm × 20 ms. In addition, we performed source reconstruction of the spatiotemporal image volumes in order to make inferences about the cortical regions that generated the scalp data. We co-registered the sensor data with a single sphere head model in order to obtain the source estimates on the individuals’ cortical mesh. Next, we conducted forward computations of the effect each dipole on the cortical mesh has on the sensors. Finally, we inverted the forward computations with the multiple sparse priors algorithm under group constraints (4, 5); these inverse reconstructions were summarized as images (smoothed at 8mm^3)^ for each of the four conditions in every participant.

For both spatiotemporal and source level, data were analysed using a mass-univariate general linear model method. We conducted a full factorial analysis, with factors: Environment (Stable and Volatile) and PE (Standards and Deviants). We computed contrast images for main effects, interactions and t-tests, in order to gauge the differential effect between deviants and standards during stable and volatile conditions. In addition, we conducted multiple regression analyses with regularity learning error as the predictor and activity at the scalp and source level as the outcomes. Age was added into all models as a covariate, since attenuation in PE response occurs with age (6). The order of volatile and stable conditions was also included as a covariate as it has been shown to influence PE responses (7, 8). Finally, psychotic experience was added as a covariate in order to exclude any potential differences in volatile and stable conditions due to psychotic symptoms. All statistical maps are reported at a threshold of *p* < 0.05 family-wise error (FWE) corrected for multiple comparisons for the spatiotemporal peak/volume or source region.

**Additional source results from the discovery dataset.** The statistical parametric maps for source-reconstructed images revealed ten significant clusters for the main effect of PE response, which overlapped with the previously reported prediction error brain network. These clusters were over bilateral inferior parietal lobule, the right motor cortex, the left middle frontal gyrus, the right occipital regions and bilateral temporal regions (see Figure S4a). In addition, the source-level multiple regression analysis at the interaction between PEs and volatility (Stable PEs > Volatile PEs), demonstrated that a decrease in regularity learning errors significantly predicted an increase in activity in the right superior frontal gyrus (peak-level *z* = 2.19, *p* uncorrected = 0.014) and right fusiform gyrus (peak-level *z* = 1.89, *p* uncorrected = 0.029); see Figure S4b.

**Table S1.** Demographic Information

|  | **Discovery dataset** | | | | **Validation dataset** | | | | ***p* value** |
| --- | --- | --- | --- | --- | --- | --- | --- | --- | --- |
|  | **Mean** | **SD** | **Min** | **Max** | **Mean** | **SD** | **Min** | **Max** |  |
| **Age (years)** | 24.65 | 4.85 | 19 | 38 | 24.27 | 5.13 | 18 | 39 | 0.75 |
| **Education (years)** | 14.29 | 2.05 | 12 | 19 | 15.66 | 3.07 | 12 | 25 | 0.034 |
| **English (years of speaking)** | 19.68 | 8.88 | 4 | 37 | 19.19 | 6.90 | 5 | 37 | 0.79 |
| **PQ score** | 3.76 | 1.21 | 0.69 | 5.73 | 4.09 | 0.80 | 2.20 | 5.57 | 0.19 |
|  | **Male** | **Female** |  |  | **Male** | **Female** |  |  |  |
| **Gender** | 14 | 17 |  |  | 22 | 22 |  |  | 0.68 |
|  | **Right** | **Left** |  |  | **Right** | **Left** |  |  |  |
| **Handedness** | 26 | 5 |  |  | 39 | 4 |  |  | 0.38 |

Abbreviations: SD = standard deviation; PQ = Prodromal questionnaire.

PQ Psychotic experience scores are log transformed.

Note: Missing information from 1 validation study participant for Education, English and Handedness.

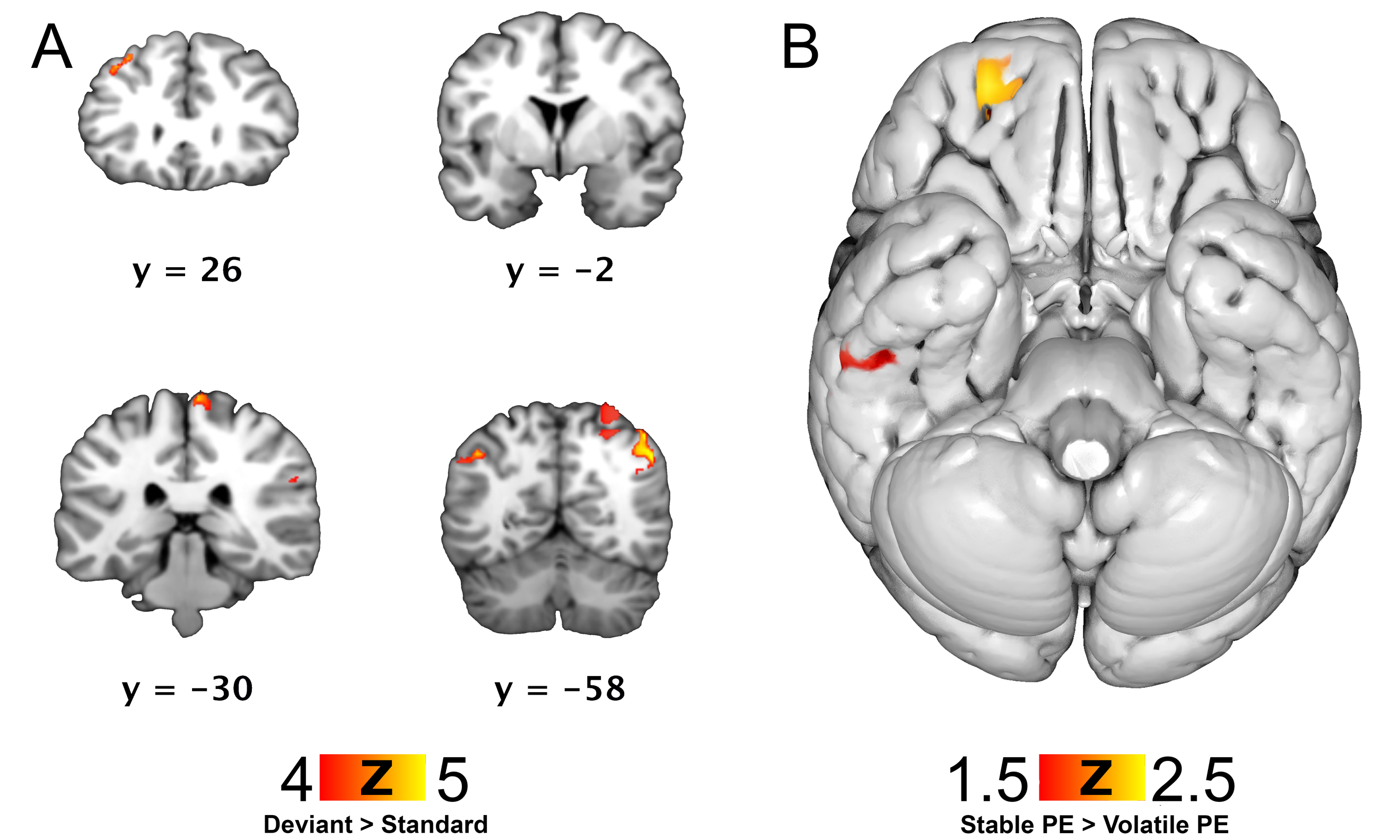

**Fig. S1. Source reconstruction analyses from the discovery dataset.** A) Main effect of prediction error response, ten clusters for deviant sounds > standard sounds; results are displayed at *p* < 0.05, FWE whole-volume corrected; B) Multiple regression analysis revealed a negative relationship between regularity learning errors and source activity in right superior frontal gyrus and right fusiform gyrus during the interaction (Stable PE > Volatile PE); results are displayed at *p* < 0.05, uncorrected.

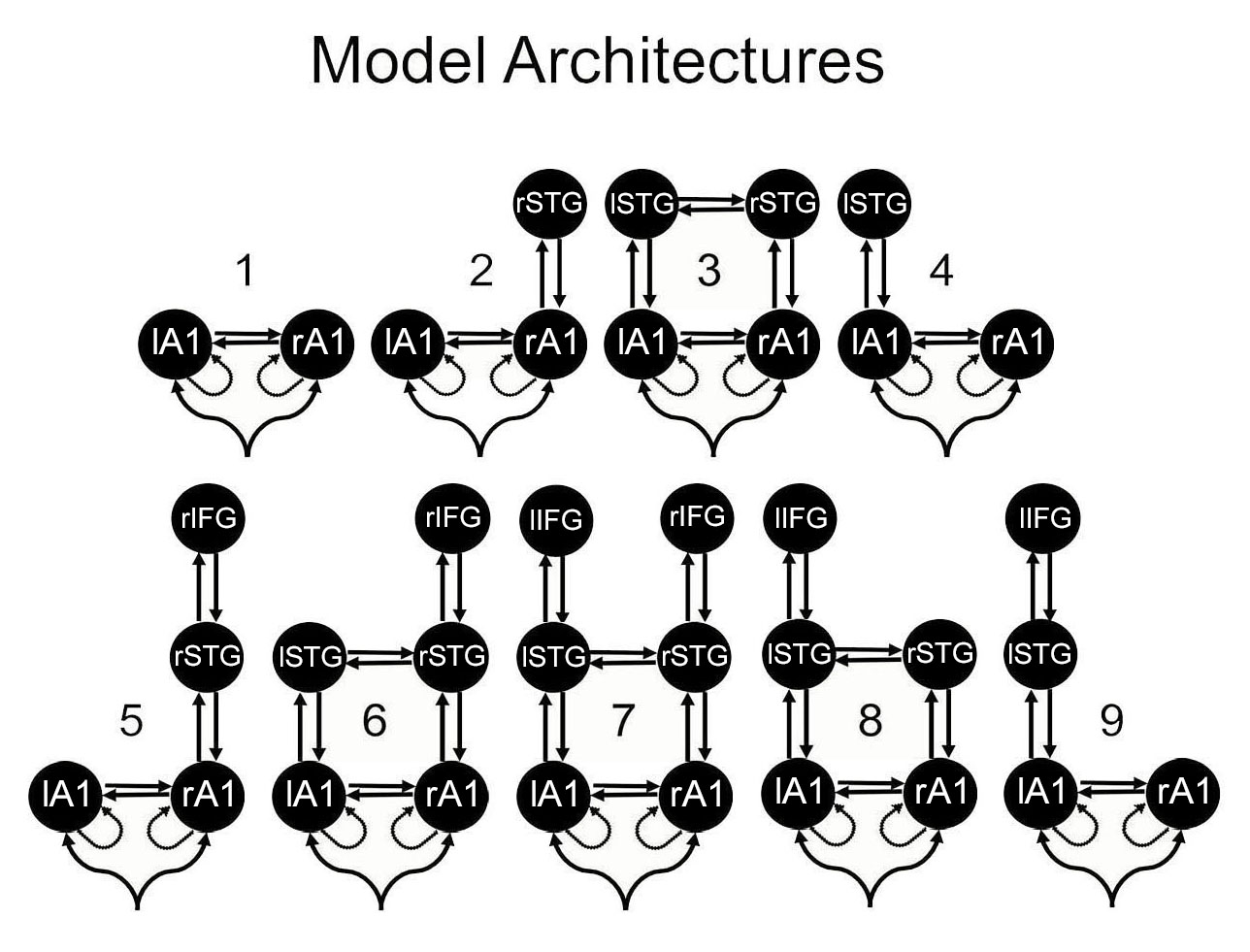

**Fig. S2. Dynamic causal models.** Nine model architectures were tested against the data. Nodes in the models included bilateral primary auditory cortices (A1), bilateral superior temporal gyri (STG), and bilateral inferior frontal gyri (IFG). [l – left, r - right]. Models with the above architectures were tested with modulations placed on 1. Forward, Backward and Intrinsic 2. Forward & Intrinsic 3. Forward & Backward 4. Null; resulting in 4 families of models and 36 models in the total model space.

**Table S2.** Pearson’s and Bayesian Correlation Matrix for Psychotic Experience, Regularity Learning and Prediction Error

|  | |  | **Regularity**  **learning error** | **Stable**  **prediction error** | **Volatile**  **prediction error** |
| --- | --- | --- | --- | --- | --- |
| ***Discovery Dataset*** | |  |  |  |  |
| **Psychotic experience** | | *r* | 0.394* | 0.326 | 0.188 |
|  |  | BF_10_ | 4.370 | 1.986 | 0.609 |
| **Regularity learning error** | | *r* | — | 0.522** | 0.327 |
|  |  | BF_10_ | — | 33.993** | 2.005 |
| ***Validation Dataset*** | |  |  |  |  |
| **Psychotic experience** | | *r* | 0.306* | -0.001 | 0.024 |
|  | | BF_10_ | 2.553 | 0.191 | 0.217 |
| **Regularity learning error** | | *r* | — | 0.324* | 0.061 |
|  | | BF_10_ | — | 3.137 | 0.267 |
|  | Table displays both Pearson’s correlations (*r*) and Bayes factors (BF_10_ ). | | | | |
|  | For Pearson’s correlations: * p < .05, ** p < .01, *** p < .001 For Bayes factors: * BF₊₀ > 10, ** , BF₊₀ > 30, *** BF₊₀ > 100 *Note*. For all tests, the alternative hypothesis specifies that the correlation is positive (_+0_). | | | | |

**Table S3.** Pearson’s and Bayesian Correlation Matrix for Psychotic Experience and Effective Connections

|  |  | **lA1** | **rA1** | **lA1** | **lSTG1** | **rA1** | **rSTG1** | **lSTG1** | **lIFG** | **rSTG1** | **rIFG** |
| --- | --- | --- | --- | --- | --- | --- | --- | --- | --- | --- | --- |
|  |  | **lA1** | **rA1** | **lSTG1** | **lIFG** | **rSTG1** | **rIFG** | **lA1** | **lSTG1** | **rA1** | **rSTG** |
| ***Discovery Dataset*** | | |  |  |  |  |  |  |  |  |  |
| **Psychotic experience** | *r* | 0.225 | -0.094 | 0.186 | -0.118 | 0.074 | 0.143 | 0.136 | 0.219 | -0.076 | -0.489** |
|  | BF_10_ | 0.108 | 0.346 | 0.12 | 0.395 | 0.169 | 0.135 | 0.138 | 0.11 | 0.316 | 18.282* |
| ***Validation Dataset*** | |  |  |  |  |  |  |  |  |  |  |
| **Psychotic experience** | *r* | 0.202 | -0.253 | -0.094 | -0.063 | 0.192 | 0.141 | 0.197 | -0.055 | -0.244 | -0.088 |
|  | BF_10_ | 0.091 | 1.258 | 0.328 | 0.271 | 0.094 | 0.110 | 0.093 | 0.260 | 1.142 | 0.316 |

Table is displaying both Pearson’s correlations (*r*) and Bayes factors (BF_10_).

For Pearson’s correlations: * p < .05, ** p < .01, *** p < .001

For Bayes factors: * BF_-_₀ > 10, ** , BF_-_₀ > 30, *** BF_-_₀ > 100

*Note*. For all tests, the alternative hypothesis specifies that the correlation is negative (_-0_).

Abbriviations: l = left; r = right; A1 = primary auditory cortex; STG = superior temporal gyrus; IFG = inferior frontal gyrus.

**Prior and Posterior	Bayes Factor Robustness Check**

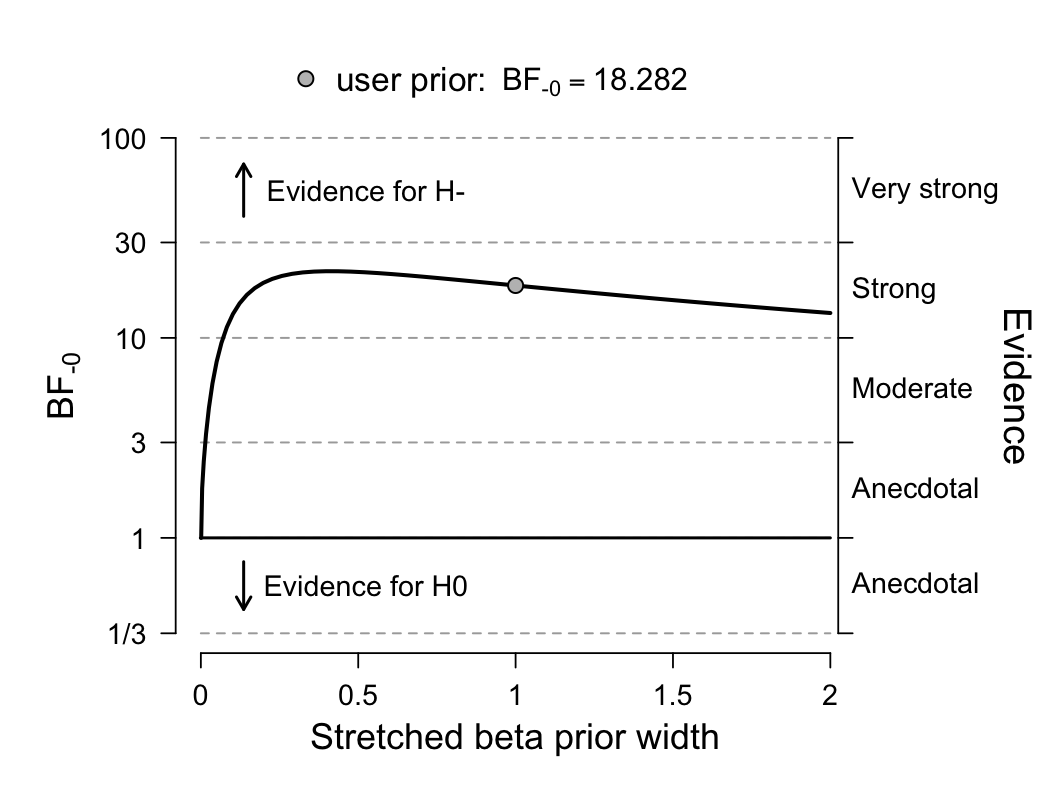

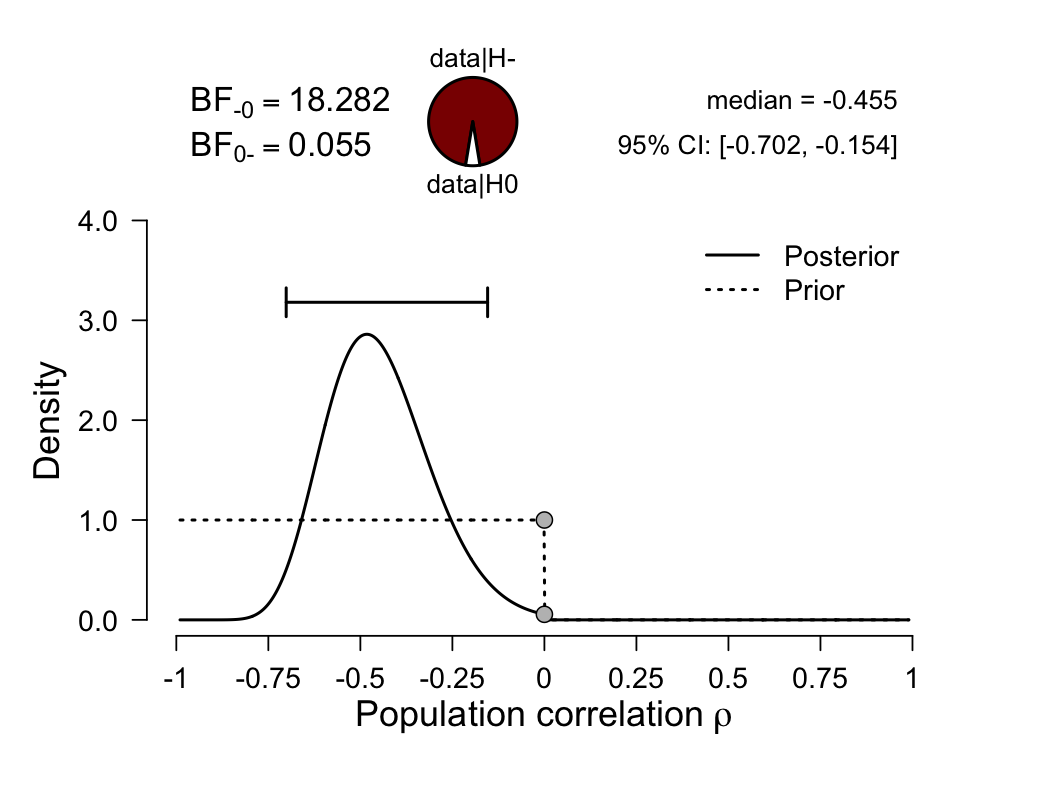

**Fig. S3. Discovery dataset prior and posterior distribution and Bayes robustness check**. This is for the correlation between psychotic experience and top-down connectivity from right inferior frontal gyrus (rIFG) to right superior temporal gyrus (rSTG). The one-sided Bayes factor, which is computed and visualized using the Savage Dickey density ratio method (9), equals 18.28 in favor of the alternative over the null hypothesis. This result indicates strong evidence for the alternative hypothesis, which is that greater psychotic experience is negatively associated with greater top-down rIFG to rSTG connectivity.
